# Structural basis of HEAT-kleisin interactions in the human condensin I subcomplex

**DOI:** 10.1101/435149

**Authors:** Kodai Hara, Kazuhisa Kinoshita, Tomoko Migita, Kei Murakami, Kenichiro Shimizu, Kozo Takeuchi, Tatsuya Hirano, Hiroshi Hashimoto

## Abstract

Condensin I is a multi-protein complex that plays an essential role in mitotic chromosome assembly and segregation in eukaryotes. It is composed of five subunits: two SMC (SMC2 and SMC4), a kleisin (CAP-H) and two HEAT-repeat (CAP-D2 and -G) subunits. Although it has been shown that balancing acts of the two HEAT-repeat subunits enable this complex to support dynamic assembly of chromosomal axes in vertebrate cells, its underlying mechanisms remain poorly understood. Here, we report the crystal structure of a human condensin I subcomplex comprising hCAP-G and hCAP-H. hCAP-H binds to the concave surfaces of a harp-shaped HEAT repeat domain of hCAP-G. A physical interaction between hCAP-G and hCAP-H is indeed essential for mitotic chromosome assembly recapitulated in *Xenopus* egg cell-free extracts. Furthermore, this study reveals that the human CAP-G-H subcomplex has the ability to interact with not only a double-stranded DNA, but also a single-stranded DNA, implicating potential, functional divergence of the vertebrate condensin I complex in mitotic chromosome assembly.

## INTRODUCTION

Immediately before cell divisions, chromatin that resides in the nucleus is converted into a set of rod-shaped structures to support their faithful segregation into daughter cells. The condensin complexes play a central role in this process, known as mitotic chromosome assembly or condensation, and also participate in diverse chromosome functions such as gene regulation, recombination and repair (Uhlmann,, 2016; Hirano, 2016). Moreover, hypomorphic mutations in the genes encoding condensin subunits have been implicated in the human disease microcephaly (Martin et al., 2016). Many eukaryotes have two different types of condensin complexes, namely, condensins I and II. Condensin I, for example, consists of a pair of SMC (structural maintenance of chromosomes) ATPase subunits (SMC2 and SMC4) and three non-SMC regulatory subunits (CAP-D2, -G and -H). SMC2 and SMC4 dimerize through their hinge domains to form a V-shaped heterodimer, and CAP-H, which belongs to the kleisin family of proteins, bridges SMC head domains through its C- and N-terminal regions. CAP-D2 and -G, both of which are composed of arrays of short amphiphilic helices, known as HEAT repeats, bind to the central region of CAP-H (Onn et al., 2007)(Yoshimura & Hirano, 2016). Although many if not all prokaryotic species have a primitive type of condensin composed of an SMC homodimer and two other regulatory subunits including a kleisin subunit, the HEAT repeat subunits are unique to eukaryotic condensins, and not found in prokaryotic condensins.

Biochemical studies using purified condensin I holocomplexes identified several ATP-dependent activities in vitro, such as positive supercoiling of DNA (Hagstrom et al., 2002; Kimura & Hirano, 1997; St-Pierre et al., 2009), DNA compaction (Strick et al., 2004), translocation along dsDNA (Terakawa et al., 2017), and DNA loop extrusion (Ganji et al., 2018). Mechanistically how these activities are supported by condensin I remains poorly understood. In fact, condensin I has the capacity to interact with DNA in many different ways. For instance, like cohesin and prokaryotic SMC complexes, it encircles double-stranded DNA (dsDNA) within its tripartite ring composed of the SMC dimer and kleisin (Cuylen et al., 2011; Ivanov & Nasmyth, 2005; Wilhelm et al., 2015). It has also been shown that a mouse SMC2-SMC4 hinge domain binds single-stranded DNA (ssDNA), but not dsDNA (Griese et al., 2010), whereas a budding yeast non-SMC subcomplex composed of YCG1/CAP-G, YCS4/CAP-D2, and BRN1/ CAP-H binds dsDNA, but not ssDNA (Piazza et al., 2014). A recent study reported the crystal structure of a budding yeast non-SMC subcomplex consisting of YCG1 and BRN1 bound to dsDNA (Kschonsak et al., 2017). Another study using *Xenopus* egg cell-free extracts showed that the pair of the HEAT repeat subunits has a critical role in dynamic assembly of mitotic chromosome axes (Kinoshita et al., 2015).

In the current study, we have determined the crystal structure of a human subcomplex composed of CAP-G bound by a short fragment of CAP-H. The structure established molecular interactions between human CAP-G and CAP-H, and implicated a role for these interactions in the ability of condensin I to support mitotic chromosome assembly. Furthermore, the human CAP-G-H subcomplex bound both dsDNA and ssDNA, implicating a potential, functional divergence of the eukaryotic condensin I complex.

## RESULTS AND DISCUSSION

### Structure of a human CAP-G-H subcomplex

The consensus sequence of HEAT repeats at the primary structure level is not tight. The original report by Neuwald and Hirano (2000) had assigned nine HEAT repeats in vertebrate CAP-G, whereas a subsequent re-assignment by Yoshimura and Hirano (2016) had identified 19 HEAT repeats that span along the near-entire length of human CAP-G (hCAP-G). Furthermore, the secondary structural prediction server PrDOS (Ishida & Kinoshita, 2007) predicted that hCAP-G has two disordered, non-HEAT regions in its central (amino-acid residues 479-553) and C-terminal (901-1015) sequences (**Fig. 1A**, **upper**). On the other hand, human CAP-H (hCAP-H) had five regions that are conserved among its orthologs among eukaryotic species (motifs I-V) (**Fig. 1A**, **lower**). Previous biochemical study had shown that the N-terminal and C-terminal halves of hCAP-H bind to hCAP-D2 and hCAP-G, respectively (Onn et al., 2007). Because the most C-terminally located motif V was predicted to bind to SMC2 (Haering et al., 2004), we thought that motif IV (residues 461-503) might be responsible for binding to hCAP-G. With these pieces of information in our hand, we aimed to express and purify hCAP-G complexed with a fragment of hCAP-H. We found that the entire HEAT domain of hCAP-G lacking the internal disordered region (residues 1-478, 554-900) and a fragment of hCAP-H containing motif IV (residues 460-515) could be co-expressed and co-purified. This hCAP-G-H subcomplex was successfully crystalized and its structure was determined at 3.0 Å resolution (**Table 1**). Two molecules of hCAP-G-H subcomplex are present in the crystallographic asymmetric unit (**Fig. EV2A**). Their structures are essentially identical, but a 4-(2-hydroxyethyl)-1-piperazineethanesulfonic acid (HEPES) is bound to only one of the two molecules. In the current report, we describe the HEPES-bound hCAP-G-H subcomplex (a, b-molecules) as a representative structure (**Fig. 1B**) in the current report.

**Table 1.** Data collection and refinement statistics

Data collection and refinement statistics
| Statistics | Native | Au (peak) |
| --- | --- | --- |
| *Data collection* |  |  |
| Wavelength (Å) | 0.98000 | 1.02991 |
| Space group | *P* 2<sub>1</sub> | *P* 2<sub>1</sub> |
| *a* (Å) | 122.4 | 123.1 |
| *b* (Å) | 62.0 | 61.7 |
| *c* (Å) | 130.9 | 131.9 |
| $\beta$ (°) | 93.4 | 93.6 |
| Resolution (Å) | 20.00 - 3.00 (3.16 - 3.00) | 20.00 - 3.38 (3.56 - 3.38) |
| Observed reflections | 132,602 (21,115) | 364,372 (54,928) |
| Unique reflections | 39,494 (6,128) | 53,098 (8,296) |
| Redundancy | 3.35 | 6.41 |
| *R*-merge | 0.069 (0.552) | 0.138 (0.863) |
| Completeness (%) | 98.6 (98.0) | 99.2 (99.0) |
| $\langle I \rangle / \sigma \langle I \rangle$ | 13.8 (2.07) | 10.26 (1.83) |
| *Refinement* |  |  |
| Resolution (Å) | 19.80 - 3.00 |  |
| Refined reflections | 39,492 |  |
| Free reflections | 1,981 |  |
| *R*-work | 0.2180 |  |
| *R*-free | 0.2732 |  |
| Root mean square deviation |  |  |
| Bond lengths (Å) | 0.003 |  |
| Bond angles (°) | 0.780 |  |
| Protein Data Bank ID | 6IGX |  |

Values in parentheses are those for the high resolution range.

**Figure 1.**
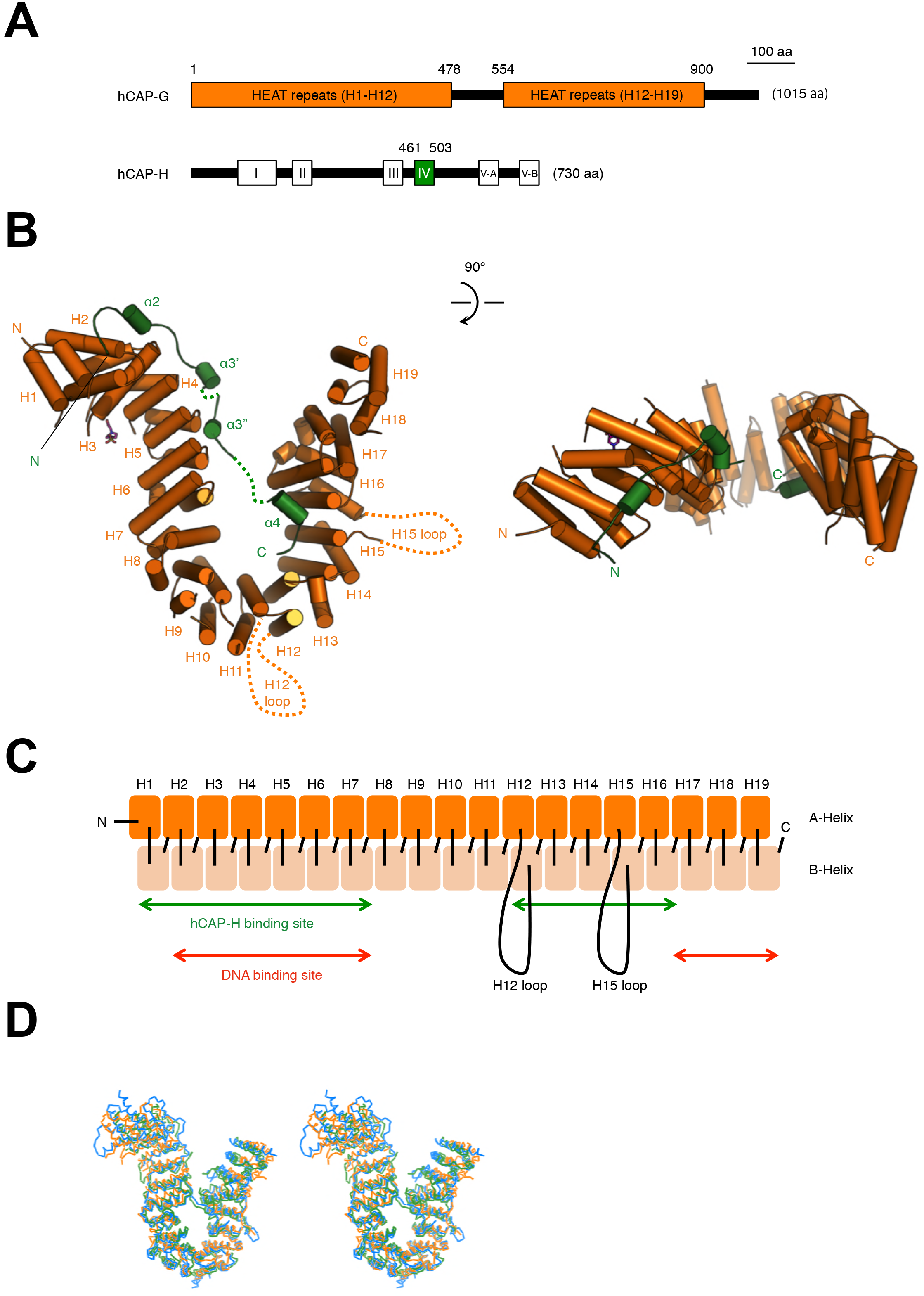
Domain architecture and overall structure of the hCAP-G-H subcomplex. **(A)** hCAP-G is 1,015 amino-acid long and contains 19 HEAT repeats. hCAP-H is 730 amino-acid long and contains 5 conserved motifs. **(B)** Cartoon diagram of the crystal structure of hCAP-G (orange) in complex with a fragment of hCAP-H (green). Unstructured, disordered regions are indicated by the dots. The 19 HEAT repeats (H1-H19) and 2 disordered loops (H12 loop and H15 loop) of hCAP-G, and 4 helices (α2, α3’, α3”, and α4) of hCAP-H are labeled. The N- and C termini of CAP-G and CAP-H are also indicated. A molecule of 4-(2-hydroxyethyl)-1-piperazineethanesulfonic acid (HEPES) is shown by the pink-colored stick model. A 90-degree rotated version is shown on the right. **(C)** Schematic illustration of the structure and domain organization of hCAP-G. Two antiparallel helices (A and B helices) comprising each HEAT repeat are colored in orange and light orange, respectively. The binding sites of hCAP-H and DNA are indicated by the green and red double-headed arrows, respectively. The H12 loop (residues 479-553) connecting the H12A and H12B helices and the H15 loop (residues 661-691) connecting the H15A and H15B helices are shown by black loops. **(D)** Comparison of the hCAP-G-H subcomplex with its related structures. Superimposition of the structures of hCAP-G-H (orange), *S. cerevisiae* YCG1-BRN1 (blue), and *S. pombe* CND3-CND2 (green) is shown as a stereo view of Cα-tracing model.

Consistent with the recent assignment based on its amino-acid sequence (Yoshimura & Hirano, 2016), hCAP-G displays a “harp-shaped” structure composed of 19 HEAT repeats (H1-H19), in which H12 and H15 have disordered loops (residues 479-553 and 661-691, respectively)(**Fig. 1, B-C**; **Fig. EV1A**; **Fig. EV2B**). hCAP-H, which comprises four α-helices (α2, α3’, α3”, and α4), binds to the concave surfaces of hCAP-G (**Fig. 1B**; **Fig. EV1B**). This overall structure in which a kleisin fragment binds to the concave surfaces of a harp-shaped HEAT repeat domain is highly reminiscent of other cohesin subunits and its regulators (Hara et al., 2014; Kikuchi et al., 2016; Ouyang et al., 2016) as well as budding and fission yeast condensin subunits (YCG1-BRN1 and CND3/CAP-G-CND2/CAP-H; Kschonsak et al., 2017). It should be noted that hCAP-G used in this study shares only 16% and 21% amino-acid identities to YCG1 and CND3, respectively, and that the hCAP-H fragment bound to hCAP-G shares only 25% and 29% identities to BRN1 and CND2, respectively. Despite the great divergence in their amino-acid sequences, two basic residues (K60, R848) located at the N- and C-terminal lobes of hCAP-G, which corresponds to K70 (YC1) and R849 (YC2) of YCG1, respectively, are conserved well (**Fig. EV1A**). Likewise, four basic residues (R435, R437, K456, and K457) of hCAP-H, which corresponds to K409 (BC1), R411 (BC1), K456 (BC2), and K457 (BC2) of BRN1, respectively, are also conserved (**Fig. EV1B**). Kschonsak et al. (2017) has recently shown that the corresponding amino-acid residues of YCG1-BRN1 contribute to dsDNA binding, strongly suggesting that the hCAP-G-H subcomplex also uses these residues to bind to dsDNA (see below).

We next performed superimpositions between the hCAP-G-H subcomplex and its budding yeast counterpart YCG1-BRN1 by using PyMoL (http://www.pymol.org/). Structural alignment between the two subcomplexes (**Fig. 1D**, **orange and blue**) shows a root mean square deviation (RMSD) value of 4.206 Å for 3,956 superimposable atoms. Structural alignment between hCAP-G-H and its fission yeast counterpart CND3-CND2 (**Fig. 1D**, **orange and green**) shows an RMSD value of 5.419 Å for 4,044 superimposable atoms. These superimpositions indicate that the overall structure of hCAP-G-H is basically identical to the structures of its yeast counterparts. There are nonetheless several notable differences between the human and yeast structures. First, some secondary structures of hCAP-G-H subcomplex are different from those of the yeast counterparts. The H12 loop is a common disordered loop also found in the yeast counterparts, but the H15 disordered loop present in hCAP-G is missing in its yeast counterparts (**Fig. 1, B-C**; **Fig. EV1A**). The hCAP-H sequence (residues 499-503), which corresponds to the buckle region of BRN1 (residues 498-504) and CND2 (residues 519-523) critical for the “safety-belt” mechanism (Kschonsak et al., 2017), is also disordered in our hCAP-G-H structure. (**Fig. EV1A**). Notably, the α3 helix of BRN1 is split into two helices (the α3’ and α3” helices) in hCAP-H, producing a disordered loop that connects the two helices (**Fig. EV1B**). Overall, the hCAP-G-H subcomplex is structurally more flexible compared with the YCG1-BRN1 subcomplex. Second, hCAP-H is more loosely bound with hCAP-G than the YCG1-BRN1 subcomplex. In fact, the distance between the NH2 of R257 and the CG2 of V754 of hCAP-G, located at H7 and H16, respectively, is 17.06 Å (**Fig. 2A**), whereas the corresponding distance between the NH2 of R287 and the CD2 of F749 of YCG1 is 6.25 Å (**Fig. EV3A**). The distance between the NZ of K154 and NZ of K889 of hCAP-G, located at H4 and H19, respectively, is 35.29 Å (**Fig. 2A**), whereas the corresponding distance between the NH1 of R170 and NZ of K895 of YCG1 is 23.02 Å (**Fig. EV3A**). These differences in the structural flexibility allow us to speculate that our hCAP-G-H structure represents an “open conformation” before binding to DNA, whereas the structure of its yeast counterpart represents a “closed conformation” after capturing dsDNA.

**Figure 2.**
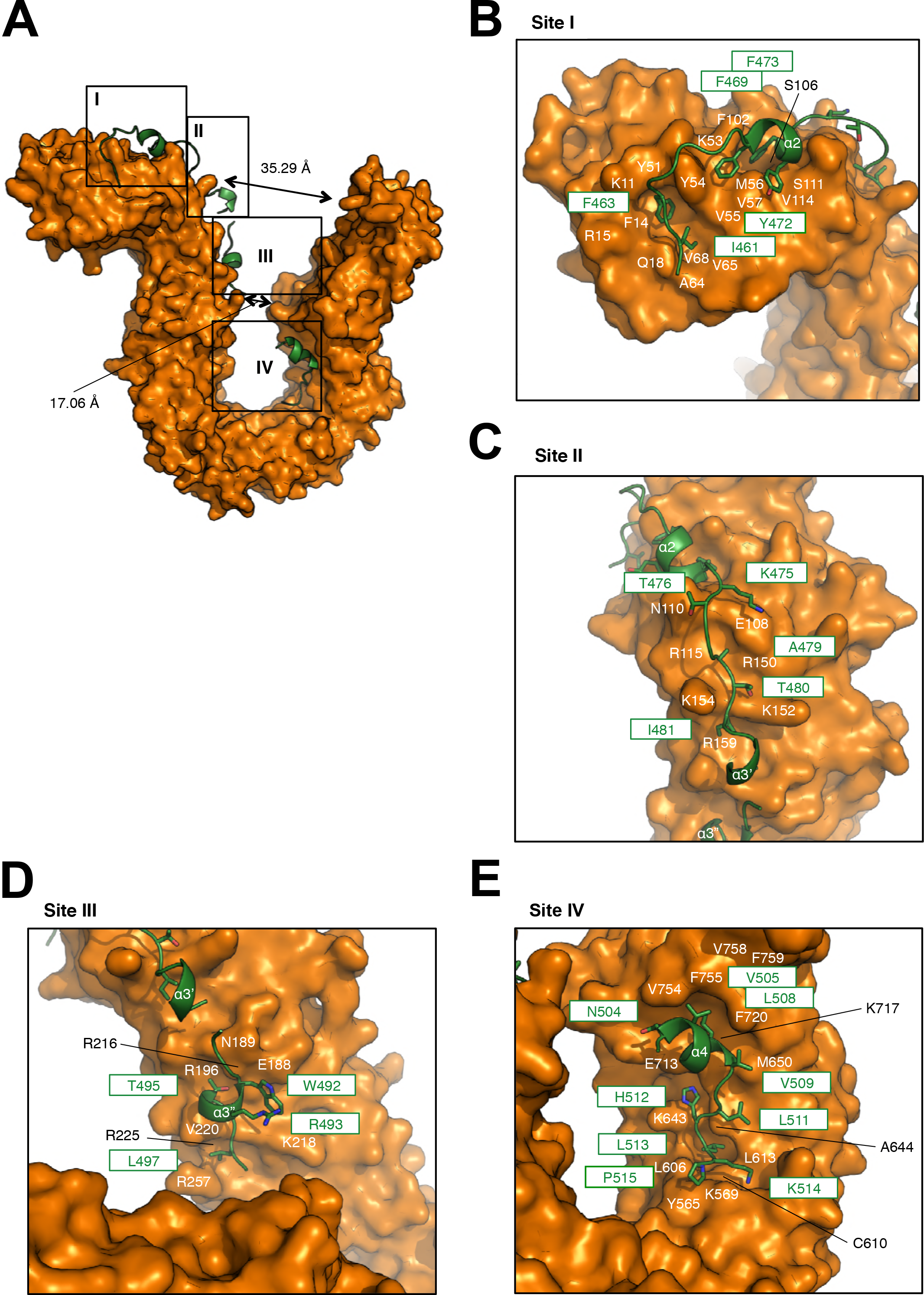
Structural details of the interaction between hCAP-G and hCAP-H. **(A)** The molecular surface of hCAP-G is shown in orange color. hCAP-H is shown as a green ribbon model. The four major contact sites (I, II, III, and IV) are boxed. **(B-E)** Zoomed-in views of sites I-IV. Residues of hCAP-G and hCAP-H are labeled in white or black and green, respectively.

### Structural details of the interaction between hCAP-G and hCAP-H

hCAP-H interacts extensively with the concave surface of hCAP-G at four major sites (**Fig. 2A**), whereas YCG1 interacts with BRN1 at five major sites (Kschonsak et al., 2017; **Fig. EV3A**). At site I, a pocket comprised of residues located at H1 (K11, F14, R15, and Q18) and H2 (Y51, Y54, V55, A64, V65, and V68) of hCAP-G accommodate I461 and F463 positioned at the N-terminal loop of hCAP-H through van der Waals contacts (**Fig. 2B**). F469, Y472, and F473 within the α2 helix of hCAP-H make mainly hydrophobic interactions with residues located at H2 (K53, M56, V57) and H3 (F102, S106, S111, and V114) of hCAP-G. In the YCG1-BRN1 subcomplex, I461, F463, E471, V474, and F475 within the N-terminal loop and the α2 helix of BRN1 form conserved hydrophobic interactions with YCG1, although van der Waals interactions also form between E460 of BRN1 and Q25 of YCG1, F464 of BRN1 and a pocket comprising F14, N15, A18, E19, and Q22 of YCG1, and T466 and N468 of BRN1 and F14 and K64 positioned at the edge of a conserved pocket of YCG1 (**Fig. EV3B**).

At site II, hCAP-H interacts with hCAP-G by van der Waals forces. K475, T476, A479, T480, and I481 of CAP-H are accommodated in a shallow pocket, where is composed of E108, N110, R115, R150, K152, K154, and R159 positioned at the H3 and H4 domains of CAP-G (**Fig. 2C**). At site II of the YCG1-BRN1 subcomplex, some residues (K478, T481, K482, I483, D484, and M485) of BRN1 also interacts with YCG1 by van der Waals interactions. Especially, I483 and M485 of BRN1 are accommodated in two deep pockets, where are composed of E127, P129, R134, R166, Y168, D169, R170, and P218 positioned at the corresponding domains of YCG1 (**Fig. EV3C**). The site II interactions in both hCAP-G-H and YCG1-BRN1 primarily involve hydrophobic interactions, but the depths of their interaction pockets are substantially different: hCAP-G recognizes the small side chain of hCAP-H, whereas YCG1 could recognizes bulky side chains of BRN1.

At site III, T495 within the α3” helix of hCAP-H is accommodated in a shallow pocket, composed of E188, N189, R196, R216, K218, and V220 positioned at the H5 and H6 domains of hCAP-G (**Fig. 2D**). A pocket comprised of residues located at H6 (V220 and R225) and H7 (R257) of hCAP-G accommodates L497 positioned at the C-terminal loop of hCAP-H through van der Waals contacts. Notably, W492 in the α3” helix forms a cation-π interaction with R493 on the same helix. This interaction may stabilize the binding of W492 of hCAP-H to E188 of hCAP-G mediated by van der Waals forces. At site III of the YCG1-BRN1 subcomplex, YCG1 interacts with BRN1 by van der Waals interactions. R490 of BRN1 forms a cation-π interaction with Y168 of YCG1, and Y496 of BRN1 forms a π-π interaction with P218 of YCG1 (**Fig. EV3D**). K491, H495, and L497 of BRN1 are accommodated in an elongated cleft, where is composed of Y168, N216, P218, R223, E242, R243, R245, and V247 positioned at the H4, H5 and H6 domains of YCG1. Site III of the YCG1-BRN1 subcomplex includes deeper clefts than that of hCAP-G, allowing YCG1 to be able to bind bulky residues of BRN1. Differences of site III could explain why amino acid sequences between hCAP-H and BRN1 are not conserved well.

At site IV, N504, V505, L508, and V509 within the α4 helix of hCAP-H are accommodated in a pocket, composed of M650, E713, K717, F720, V754, F755, V758, and F759 located at the H14, H15, and H16 domains of hCAP-G (**Fig. 2E**). Especially, the conserved L508 of hCAP-H contributes to anchor the α4 helix to a hydrophobic cleft. Five residues (L511, H512, L513, K514, and P515) located at the C-terminal loop of hCAP-H are also accommodated in an elongated cleft, composed of Y565, K569, L606, C610, L613, K643, and A644 located at the H12, H13, and H14 domains of hCAP-G. H512 and L513 of hCAP-H look like “the plug”, which puts in the outlet formed by a cleft of hCAP-G. I509, F513, and I514 of BRN1 corresponding to L508, H512, and L513 of hCAP-H also work as a fine tuner and the plug for hydrophobic interactions with YCG1, respectively (**Fig. EV3F**). At site IV, there are also notable hydrogen bonds formed between L511 and H512 of hCAP-H and D647 of hCAP-G (**Fig. 3B**). D647 is an acidic residue broadly conserved among the CAP-G/YCG1 orthologs (**Fig. EV1A**). Interestingly, an aspartate side chain that makes hydrogen bonds with two backbone amides of residues in a pocket is commonly found in the prefusion state of hemagglutinin (HA) of the influenza virus (Ivanovic, Choi, Whelan, van Oijen, & Harrison, 2013) and a binding hotspot between cohesin’s HEAT repeat subunit SA2 and kleisin subunit Scc1 (Hara et al., 2014).

**Figure 3.**
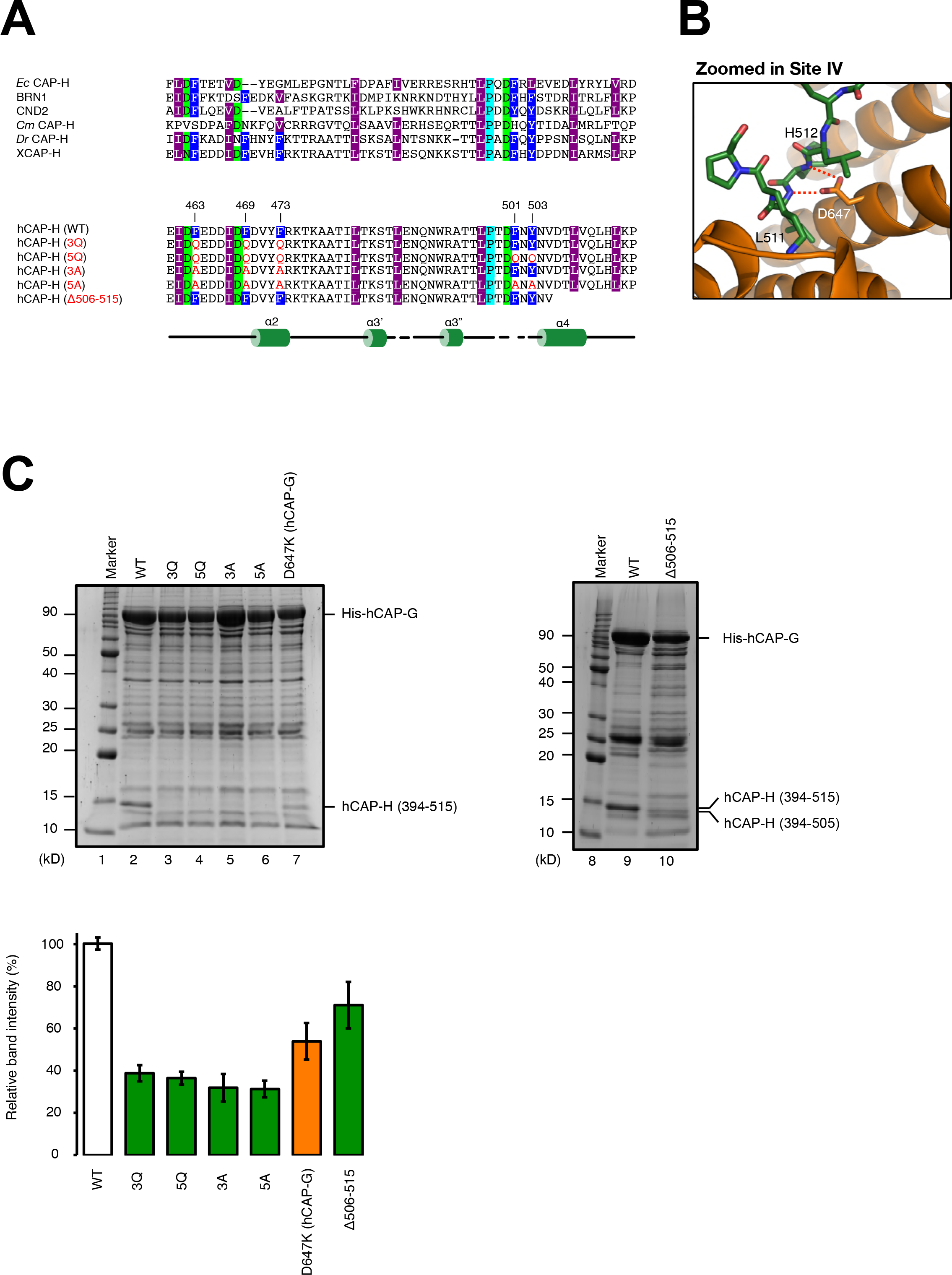
Identification of residues critical for the interaction between hCAP-G and hCAP-H. **(A)** IV-3Q, 5Q, 3A, 5A, and ∆506-515 mutants of hCAP-H. Motif IV (residues 461-503) contains amino-acid residues highly conserved among eukaryotic species (X, *Xenopus laevis*; Dr, *Danio rerio*; Cm, *Cyanidioschyzon merolae*; Sp, *Schyzosaccharomyces pombe*; Sc, *Saccharomyces cerevisiae*; Ec, *Encephalitozoon cuniculi*). To produce the IV-3Q, 5Q, 3A, and 5A mutants, the conserved aromatic amino-acid residues (F463, F469, F473, F501 and Y503; labeled in dark blue) were substituted with glutamine (Q) or alanine (A) residues. The secondary structural elements of hCAP-H is drawn below the sequence alignments. **(B)** Zoomed-in view of site IV. Residues of hCAP-G and hCAP-H are shown in orange and green, respectively. The dashed red lines indicate hydrogen bonds. **(C)** Interaction analysis between hCAP-G and hCAP-H. Bacterial cell lysates co-expressing hCAP-G (residues 1-478, 554-900) and hCAP-H (residues 394-515), either wild type (WT; lane 2, and 8), 3Q (F463Q, F469Q, and F473Q; lane 3), 5Q (F463Q, F469Q, F473Q, F501Q, and Y503Q; lane 4), 3A (F463A, F469A, and F473A; lane 5), or 5A (F463A, F469A, F473A, F501A, and Y503A; lane 6), a C-terminal deletion (506-514 residues were deleted from 394-515; lane 10), were applied to Ni-NTA agarose resin, and the bound fraction were analyzed by SDS-PAGE. Alternatively, a cell lysate co-expressing a mutant hCAP-G (D647K) and wild-type hCAP-H was tested (lane 7). The relative band intensities of hCAP-H divided by the band intensities of hCAP-G were normalized to those of the wild type (WT), and plotted with standard error (S.E.) bars (n = 3) on the lower panel.

The YCG1-BRN1 subcomplex has an additional HEAT-kleisin interaction site, site III’ (**Fig. EV3E**). At site III’, L498, P499, D501, F502, and F504 of BRN1 interact with a crack between two cliffs formed by R252, R245, V247, R287, E708, E741, A742, Q745, A746, F749, and E741 positioned at the H6, H7, and H16 domains of YCG1 (**Fig. EV3E**). Although no interactions corresponding to site III’ are found in the hCAP-G-H subcomplex, F501 and Y503 of hCAP-H corresponding to F502 and F504 of BRN1 are highly conserved among eukaryotic species. It is therefore possible that the hCAP-G-H subcomplex undergoes conformational changes (from an open form to a closed form) upon binding to dsDNA, forming site III’ interactions found in the YCG1-BRN1 subcomplex.

### Identification of residues critical for the interaction between hCAP-G and hCAP-H

To identify residues critical for the interaction between hCAP-G and hCAP-H, we designed six mutants that target conserved, surface-exposed residues at the hCAP-G-H interface, and the amount of hCAP-H fragment that co-purified with immobilized His_6_-tagged hCAP-G was evaluated (**Fig. 3, A & C)**. As expected, three Gln substitutions (3Q) of F463, F469, and F473 of hCAP-H positioned at site I greatly impaired the interaction between hCAP-G and hCAP-H (**Fig. 3C**, **lanes 2 and 3**). Three Ala substitutions (3A) of the same residues also diminished the interaction, suggesting that van der Waals interactions formed by these aromatic residues of CAP-H are crucial for its interaction with hCAP-G (**Fig. 3C, lane 5**). More interestingly, additional Gln or Ala substitutions (5Q or 5A) of F501 and Y503 of hCAP-H positioned at site III’ did not further impaired its interaction with hCAP-G (**Fig. 3C**, **lanes 4 and 6**).

These results suggest that F501 and Y503 are not directly involved in hCAP-G-H subcomplex formation, but may be required for the stabilization of a closed conformation after dsDNA binding. A Lys substitution of D647 of hCAP-G positioned at site IV (D647K) decreased its interaction with hCAP-H, suggesting that D647-mediated hydrogen bonds with L511 and H512 of hCAP-H are crucial for its interaction with hCAP-G at site IV (**Fig. 3B**; **Fig. 3C**, **lane 7**). We also found that deletion of a C-terminal region of hCAP-H (506-514 residues) also reduced its interaction with hCAP-G (**Fig. 3C**, **lanes 9 and 10**), supporting the idea that plug-outlet interactions formed by the site IV are critical for hCAP-G-H subcomplex formation, just like site I.

### The interaction between hCAP-G and hCAP-H is essential for proper chromosome assembly mediated by condensin I in Xenopus egg extracts

To test whether the interaction between hCAP-G and hCAP-H is indeed essential for the function of condensin I, we introduced the motif IV quintuple mutations (F463Q, F469Q, F473Q, F501Q, Y503Q; designated IV-5Q) described above into the context of full-length, holocomplexes **(Fig. 3A)**. By using the baculovirus expression system described previously (Kinoshita et al., 2015), we co-expressed the five subunits of mammalian condensin I containing either the wild-type or mutant form of hCAP-H in insect cells. An equal level of expression of the five subunits in the two samples was confirmed by immunoblotting against total lysates (**Fig. 4A**). Both lysates were then subjected to affinity-purification using glutathione-agarose beads (Note that the SMC4 subunit was GST-tagged), followed by proteolytic cleavage of the GST moiety. Although wild-type hCAP-G was successfully co-purified along with the other four subunits, the IV-5Q mutant form of hCAP-G was almost completely missing from the purified fraction (**Fig. 4B**). The complexes purified from the wild-type and mutant lysates were then added back into *Xenopus* egg extracts depleted of endogenous condensins (Kinoshita et al., 2015). We found that, although the holocomplex purified from the wild-type lysate produced normal chromosomes (**Fig. 4C**, **top panels**), the complex purified from the mutant lysate failed to do so, making abnormal chromosomes with fuzzy surfaces and thin axes (**Fig. 4C**, **middle panels**). The abnormal structure was highly reminiscent of those produced by the tetrameric mutant complex that lacks the hCAP-G subunit (i.e., ΔG) we reported previously (**Fig. 4C**, **bottom panels**). These results strongly suggest that the IV-5Q mutations disrupt both physical and functional interactions between hCAP-G and hCAP-H, resulting in the formation of a tetrameric mutant complex that is equivalent to the ΔG complex.

**Figure 4.**
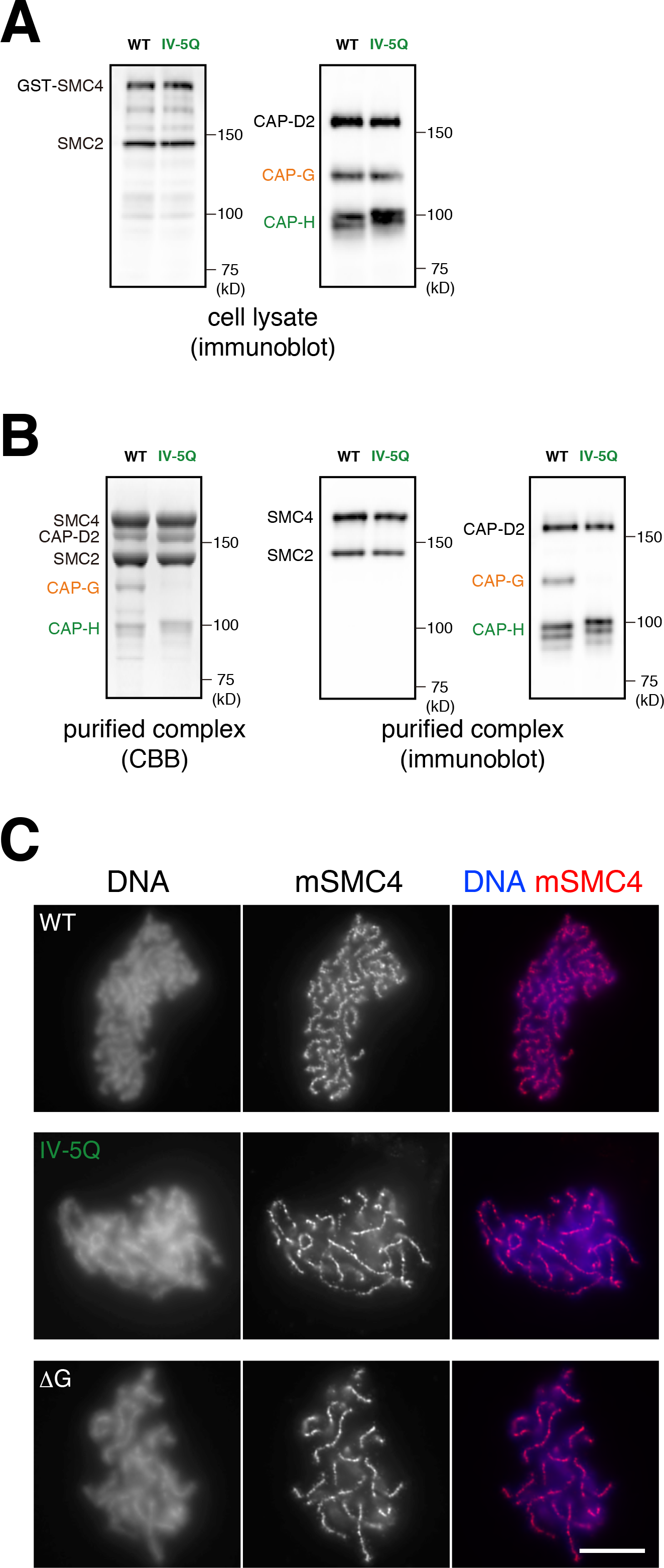
The hCAP-G-H interaction is essential for chromosome assembly in *Xenopus* egg extracts. **(A)** Expression of condensin I subunits in insect cells. The wild-type (WT) or IV-5Q mutant CAP-H subunit was co-expressed with the other four subunits (GST-SMC4, SMC2, CAP-D2 and CAP-G) in insect cells. Cell lysates were prepared and subjected to SDS-PAGE, followed by immunoblotting with a mixture of antibodies against SMC2 and SMC4 (left panel) or against CAP-D2, -G, and -H (right panel). **(B)** Purification of the WT and IV-5Q mutant condensin I complexes. Protein samples purified through glutathione-affinity chromatography were subjected to SDS-PAGE and analyzed by CBB staining (left panel) or immunoblotting with a mixture of antibodies as described above (middle and right panels). **(C)** Add-back assay using the WT and mutant condensin I complexes. *Xenopus* egg extracts depleted of endogenous condensin complexes were supplemented with the purified complexes (top, WT; middle, IV-5Q: bottom, ΔG). The supplemented extracts were then incubated with sperm nuclei to assemble mitotic chromosomes. The samples were fixed and labeled with an antibody against mSMC4 (red). DNA was counterstained with DAPI (blue). The data from a single representative experiment out of two repeats are shown. In the experiment shown here, multiple images were collected for condensin-depleted extracts supplemented with the WT (n = 17), IV-5Q (n = 22) and ΔG (n = 20) complexes. Chromosome structures produced in each extract are highly homogenous, readily allowing us to assign unique phenotypes to different conditions. The scale bar represents 10 μm.

### Identification of DNA-binding site in the hCAP-G-H subcomplex

Kschonsak et al. (2017) reported the structure of a YCG1-BRN1-dsDNA ternary complex. Despite numerous trials, however, we could not get any crystals of the corresponding ternary complex using hCAP-G and hCAP-H. To test whether our hCAP-G-H subcomplex has the ability to interact with DNA, we performed electrophoretic mobility shift assay (EMSA) using a blunt-ended dsDNA probe and a ssDNA probe. The result clearly showed that the hCAP-G-H subcomplex interacts not only with dsDNA (**Fig. 5, A & B**, **lanes 1-4**) but also with ssDNA (**Fig. 5C**, **lanes 1-4)**. To clarify the crucial residues that interacted with DNA, we mapped potential DNA-binding residues on hCAP-G-H subcomplex using structural information from the YCG1-BRN1-dsDNA ternary complex (**Fig. 5D**). It was estimated that the DNA-binding interface of the hCAP-G-H subcomplex is basically similar to that of its budding yeast counterpart. We picked up two positively charged residues (K60 and R848), which correspond to the DNA-binding residues of YCG1 (YC1/2), and constructed a K60D/R848E double mutant. The K60D/R848E double mutation greatly impaired the binding affinities to both dsDNA and ssDNA (**Fig. 5, A & B-C**, **lanes 5-8**). The results suggested that dsDNA and ssDNA binding interfaces are overlapped, and that the N- and C-terminal HEAT-repeat domains of hCAP-G are likely to be required for binding to both DNA substrates. Next, we mutated R168 of hCAP-G, a residue that is important for HEPES binding (**Fig. 5E**) and potentially confers DNA binding as well. We found that a R168E mutant reduced, but not completely eliminated, the affinities for both dsDNA and ssDNA (Fig. 5, A & B-C, **lanes 9-12**). Because the small concave surface containing R168 has no enough space to accommodate dsDNA, we speculate that it may adapt open-mouth structure to grab ds DNA. Alternatively, a conformational change of this surface could indirectly affect the DNA binding domain constituted by K60 and R848.

**Figure 5.**
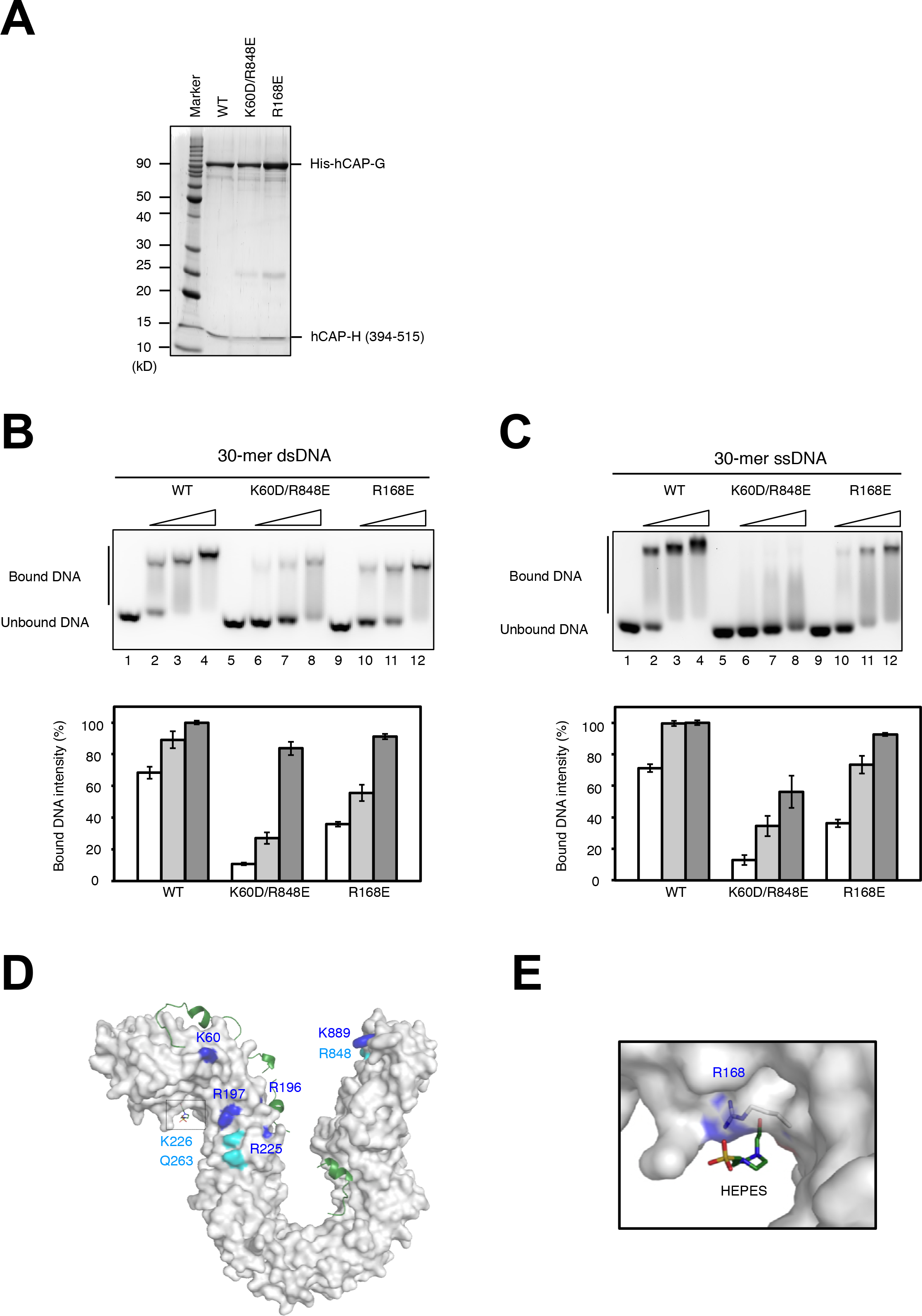
DNA-binding surfaces conserved between hCAP-G and YCG1. **(A)** Purification of hCAP-G-H subcomplexes: wild type (WT), CAP-G K60D/R848E double mutant (K60D/R848E), and CAP-G R168E mutant (R168E). Purified protein samples were subjected to SDS-PAGE and analyzed by CBB staining. **(B)** Double-stranded DNA (dsDNA) binding assay. 30-bp dsDNA were incubated with no protein (lanes 1, 5, and 9), increasing amounts of the hCAP-G-H subcomplex wild type (WT; lanes 2-4), CAP-G K60D/R848E double mutant (K60D/R848E; lanes 6-8), and CAP-G R168E mutant (R168E; lanes 10-12) (upper panel). The bound DNA intensities divided by the total DNA intensities were normalized to those of the wild type (WT). Values of the relative band intensities are visualized as bar graphs with standard error (S.E.) bars (n = 3) in the lower panel. **(C)** Single-stranded DNA (ssDNA) binding assay of the hCAP-G-H subcomplexes used in panel (**B**). **(D)** The molecular surface of hCAP-G in complex with hCAP-H. The structural model of hCAP-G is shown in white color. hCAP-H is shown as a green ribbon model. Identical and homologous residues between hCAP-G and YCG1 are shown in blue and cyan, respectively. **(E)** Zoomed-in view of the HEPES-binding site. R168 of hCAP-H interacts with HEPES.

In this study, we have determined the crystal structure of a hCAP-G-H subcomplex, in which the kleisin subunit hCAP-H is more loosely bound to the HEAT subunit hCAP-G, compared with the yeast YCG1-BRN1 complex. We have also demonstrated the structural basis of the interaction between hCAP-G and hCAP-H, whereby hCAP-H binds to hCAP-G with two conserved N- and C-terminal concave surfaces. Despite their great sequence divergences, the human and yeast structures are surprisingly similar to each other, implicating that the basic mechanisms of condensin-mediated chromosome condensation must be widely conserved among eukaryotic species with large and small chromosomes. Our functional assay employing *Xenopus* egg extracts has convincingly shown that the hCAP-G-H interaction is indeed essential for proper mitotic chromosome assembly. It should be noted that the two different mutant complexes lacking hCAP-G (IV-5Q and ΔG) still retain the ability to be loaded onto chromosomes in our cell-free assay, a result contrary to the prediction from a previous study (Kschonsak et al., 2017). On the other hand, the ability of the hCAP-G-H subcomplex to interact with ssDNA, which had not been shown for its yeast counterparts, could be related with the finding that condensin binds ssDNA segments in transcribing regions to regenerate dsDNA (Sutani et al., 2015). It will be very important in the future to further clarify the similarities and detailed differences in the structure and function of this fundamental chromosome organizing machine among different eukaryotic species.

## MATERIALS AND METHODS

### Protein production and purification

cDNAs corresponding to hCAP-G (amino-acid residues 1-900) and hCAP-H (amino-acid residues 460-515) were cloned into *Bam*HI-*Hind*III and *Nde*I-*Xho*I sites, respectively, of a pETDuet-1 vector (Novagen). Based on the result of a secondary structural prediction, we deleted a putative disordered region of CAP-G (residues 479-553) by PCR-based mutagenesis. The final constructs, which encoded an N-terminally His_6_-tagged hCAP-G (residues 1-478, 554-900) and a part of hCAP-H (460-515), was used to transform *Escherichia coli* BL21 (DE3). Cells were grown at 37C to a cell density of about 0.8 at 660 nm in LB medium, and then cultured for a further ~20 hours at 25C after the addition of 0.2 mM isopropyl β-D-1-thiogalactopyranoside (IPTG). The cells were harvested, resuspended in 10 mL of buffer I (50 mM HEPES-NaOH pH 6.8 and 250 mM NaCl) per gram of cells, and lysed by sonication. The cell lysate was clarified by centrifugation for 1 hour at 4C (48,300x *g*). The supernatant was applied to a 5-mL HiTrap Heparin HP column (GE Healthcare), and the bound proteins were eluted with a linear gradient of 250 to 800 mM NaCl over a total volume of 95 mL. The collected proteins were diluted with buffer II (50 mM Tris-HCl pH 8.5), and applied to a 5-mL HiTrap Q HP anion-exchange column (GE Healthcare). The bound proteins were eluted with a linear gradient of 0 to 600 mM NaCl over a total volume of 95 mL. The eluted proteins were passed through a HiLoad 16/600 Superdex 200 size-exclusion column (GE Healthcare) equilibrated with buffer III (20 mM HEPES-NaOH pH 7.4, 100 mM NaCl, and 5 mM DTT), and then concentrated to 15 mg/mL using a Vivaspin (30 kDa MWCO) concentrator (Sartorius). The purity of the hCAP-G-H subcomplex was confirmed by SDS-PAGE followed by Coomassie Brilliant Blue (CBB) staining. The purified protein was frozen with liquid N_2_ and stored at −80C until use.

### Crystallization, data collection, and structure determination

Crystallization of the hCAP-G-H subcomplex was performed by the sitting drop vapor-diffusion method using a commercial kit from Hampton Research, Qiagen and Molecular Dimensions to screen crystallization conditions. Drops were prepared by mixing 0.5 μL of protein solution with 0.5 μL of reservoir solution. Crystals were obtained in a few conditions with polyethylene glycol as a precipitant after a week at 20C. Conditions were further optimized with the hanging-drop vapor diffusion method. hCAP-G-H subcomplex crystals suitable for X-ray diffraction experiments appeared within 1 week with a reservoir solution consisting of 6.5% (w/v) PEG3350, 0.10 M MgCl_2_, 0.10 M HEPES-NaOH pH7.5, and 3% (v/v) ethylene glycol. Heavy atom derivatives of crystals were prepared by the soaking method using a solution of 1 mM Potassium dicyanoaurate (I), 7-12% (w/v) PEG3350, 0.10 M MgCl_2_, and 0.10 M HEPES-NaOH pH7.5 for 10 min. All crystals were cryoprotected with a reservoir solution including 20-25% (v/v) ethylene glycol before being flash-frozen.

Each crystal was picked up in a nylon loop, and cooled and stored in liquid N_2_ gas *via* an Universal V1-Puck (Crystal Positioning System Inc.) until use. X-ray diffraction data of frozen crystals were collected under a stream of N_2_ gas at −173C on the BL-17A beamline at Photon Factory (Tsukuba, Japan) using a pixel array photon-counting detector, PILATUS3 S6M (DECTRIS). The hCAP-G-H subcomplex crystal diffracted to 3.0 Å. Diffraction data were integrated, scaled, and averaged with program *XDS* (Kabsch, 2010) and *SCALA* (Evans, 2006).

Initial phases for the Au-labeled hCAP-G-H subcomplex was obtained by the single-wavelength anomalous dispersion (SAD) with AutoSol in the *PHENIX* package (Adams et al., 2010). Model building of the hCAP-G N-terminal and C-terminal HEAT repeats and hCAP-H was done with AutoBuild in *PHENIX*. Subsequent model building, especially HEAT repeats of middle region of the hCAP-G was carried out with COOT (Emsley & Cowtan, 2004), and the structure was refined with *PHENIX.REFINE*. The data collection and refinement statistics were summarized in **Table 1**. All structure drawings in this study were created with PyMOL (http://www.pymol.org/) and depicted a-molecule and b-molecule as a representative structure.

### Interaction analysis of hCAP-G and hCAP-H co-expressed in E. coli

A cDNA encoding a central part of hCAP-H (residues 394-515) was cloned into the *Nde*I-*Xho*I site of pETDuet-1 containing the cDNA of hCAP-G (residues 1-478, 554-900) in the *Bam*HI-*Hind*III site. Point mutations in the hCAP-G or hCAP-H sequence were introduced by using a PCR-based method. His_6_-tagged hCAP-G was co-expressed with the hCAP-H by a procedure similar to that described above, except that bacterial cells were incubated at 15C or 25C after IPTG induction. Interaction analysis, based on immobilized metal affinity chromatography (IMAC), was performed. In brief, cell lysates were applied to Ni-NTA agarose resin (Qiagen), and the beads were washed first with buffer IV (50 mM HEPES-NaOH pH 7.4, 1.5 M NaCl, and 20 mM imidazole) and then with buffer V (50 mM HEPES-NaOH pH 7.4 and 100 mM NaCl). The bound proteins were subjected to SDS-PAGE followed by CBB staining, and quantified using a ChemiDoc Touch Imaging System (Bio-Rad Laboratories).

### DNA-binding assay

To obtain proteins that are used for EMSA, mutations were introduced by the same method as described above. Mutants were overexpressed and purified with Ni-NTA agarose resin. The bound protein was washed with buffer IV and buffer V, and then eluted with a stepwise gradient from 50 to 500 mM imidazole. The eluted proteins were further purified by HiTrap Q and HiLoad Superdex 200. Purified mutant proteins were concentrated, frozen with liquid N_2_, and stored at −80C until use.

To investigate the preference of the hCAP-G-H subcomplex for DNA structures, EMSA was performed using 30-mer ssDNA (5’-CCTATAGTGAGTCGTATTACAATTCACTCG-3’) and 30-mer blunt-ended dsDNAs (5’-CCTATAGTGAGTCGTATTACAATTCACTCG-3’; 5’- CGAGTGAATTGTAATACGACT CACTATAGG-3’). The DNA and the subcomplex were mixed at 1:1, 1:2, and 1:4 molar ratio and incubated for overnight at 4C. Final concentration of DNA after mixing solutions was 6.7 μM. These solutions were separated by electrophoresis at 4C on 1% agarose gel containing GelRed DNA stain (Biotium) and bands were detected by a ChemiDoc Touch Imaging System. EMSA was performed at least three times and the band intensities were measured using the program ImageJ (NIH, USA). The relative band intensity was calculated by dividing the intensity of bound DNA band by the total intensity in the lane and normalized by the values of 26.8 μM wild type.

### Expression and purification of recombinant condensin complexes

To construct the IV-5Q mutant of hCAP-H, we used QuikChange Site-Directed Mutagenesis Kit (Agilent Technologies) to introduce a set of point mutations sequentially into the original expression construct (pFH101; Onn et al., 2007) so that five amino-acids (F463, F469, F473, F501 and Y503) in its coding sequence were substituted with glutamine (Q). Oligonucleotides used for mutagenesis were as follows (mutation sites introduced were underlined): F469Q, 5’-GAAGATGATATTGAC<u>CAA</u>GATGTATATTTTAGA-3’; F501Q Y503Q, 5’-CCTTCCTACAGAT<u>CAA</u>AAC<u>CAG</u>AATGTTGACACTCT-3’; F463Q, 5’- GATTTTGAAATTGAC<u>CAA</u>GAAGATGATATTGAC-3’; F469Q F473Q, 5’- GAC<u>CAA</u>GATGTATAT<u>CAA</u>AGAAAAACAAAGGCT-3’. The resultant construct (pHM110) was used for the preparation of bacmid DNA to produce a baculovirus. Expression of condensin holocomplexes and subcomplexes in insect cells and their purification were performed as described previously (Kinoshita et al., 2015).

### Chromosome assembly assays and immunofluorescence analyses

Chromosome assembly assays using *Xenopus* egg extracts and immunofluorescence analyses of chromosomes assembled in the extracts were performed as described previously (Kinoshita et al., 2015).

## ACKNOWLEDGMENT

We acknowledge the kind support of the beamline staff of Photon Factory for data collection. This work was supported by Grant-in-Aid for Scientific Research, KAKENHI [grant nos. 15K18491 and 17K07314 to KH, 15K06959 to KK, 16H04755 and 17H06014 to HH, and 15H05971 to TH] and grants from the Takeda Science Foundation and Naito Foundation (HH).

## AUTHOR CONTRIBUTION

KH and TH designed the experiments. KH, TM, and KS carried out recombinant protein production, crystallization, and structural determination. KH and KM carried out *in vitro* interaction assay, and DNA binding assay. KT performed mutagenesis of hCAP-H, and KK performed the purification of the recombinant holocomplexes and functional assays using *Xenopus* egg cell-free extracts. KH, TH and HH wrote the manuscript.

## CONFLICT OF INTEREST

The authors declare that they have no conflicts of interest.

## Expanded View Figure legends

**Figure EV1.**
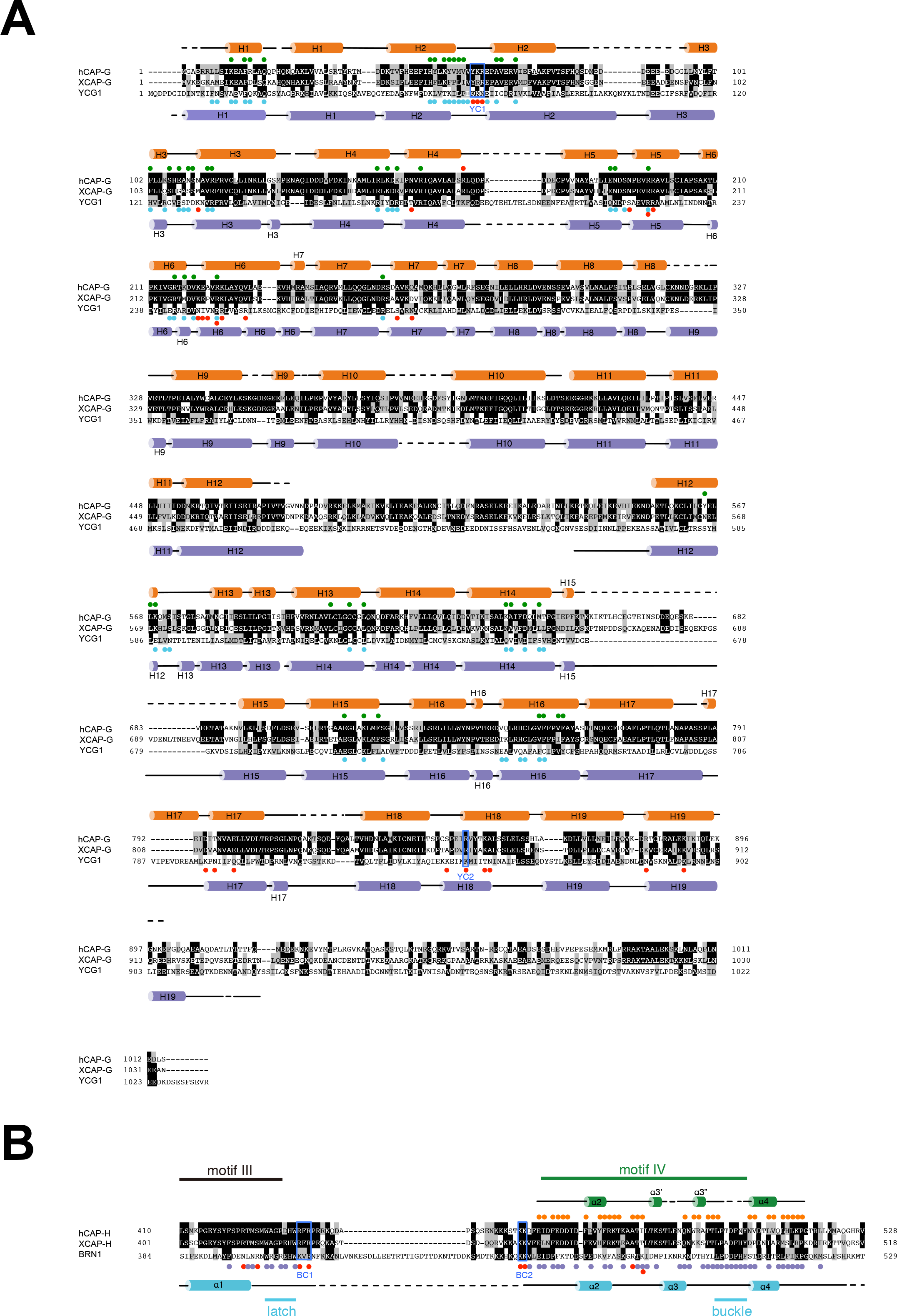
**(A)** Secondary structures and structure-based sequence alignment of human CAP-G (hCAP-G), *Xenopus* laevis CAP-G (XCAP-G), and *Saccharomyces cerevisiae* YCG1. The secondary structural elements of hCAP-G and YCG1 are drawn above and below the sequence alignments, respectively. Identical and homologous residues are shown on black and gray backgrounds, respectively. The colored circles indicate residues of hCAP-G that interact with hCAP-H (green), bind to HEPES (red). Residues of YCG1 that interact with BRN1 and dsDNA are labeled with light blue and red color, respectively. The YC1 and YC2 regions indicate residues critical for DNA binding defined by Kschonsak et al (2017). **(B)** Structure-based sequence alignment of hCAP-H, XCAP-H, and BRN1. The secondary structural elements of hCAP-H and BRN1 are drawn above and below the sequence alignments, respectively. Identical and homologous residues are shown on black and gray backgrounds, respectively. The colored circles indicate residues of hCAP-H that interact with hCAP-G (orange), and residues of BRN1 that interact with YCG1 (purple) or dsDNA (red). Also indicated are BC1, BC2, latch, and buckle regions defined by Kschonsak et al (2017), and motif III and IV of CAP-H shown in Figure 1A.

**Figure EV2.**
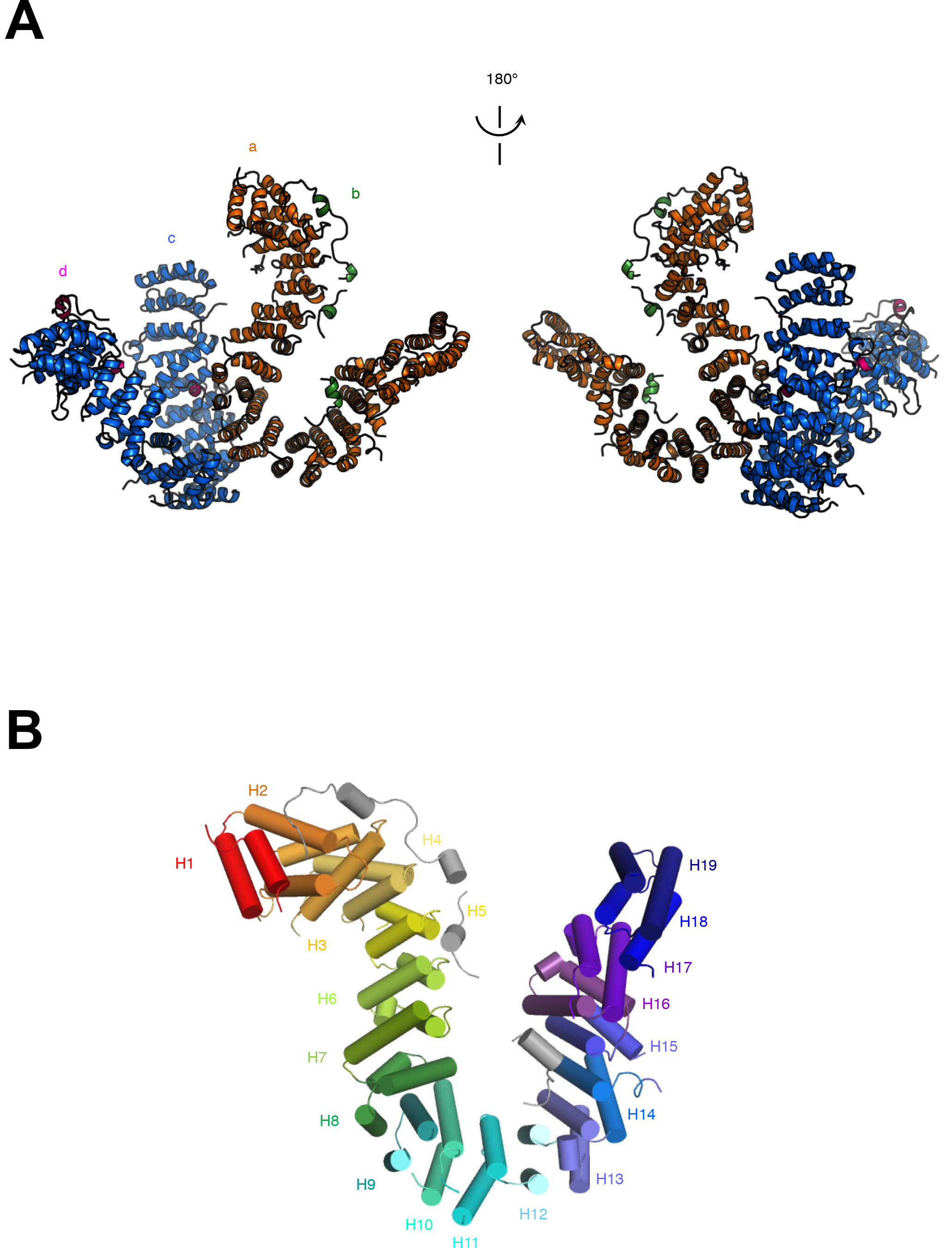
Structure of the hCAP-G-H subcomplex. **(A)** Two molecules of the hCAP-G-H subcomplex in the asymmetric unit, shown by orange (hCAP-G; a) and green (hCAP-H; b), and blue (hCAP-G; c) and pink (hCAP-H; d) ribbon representations. The pink stick model indicates HEPES. Note that HEPES bound only one of the two hCAP-G molecules (a-molecule) present in the asymmetric unit. **(B)** Cartoon representation of the 19 HEAT repeats present in hCAP-G complexed with hCAP-H (gray).

**Figure EV3.**
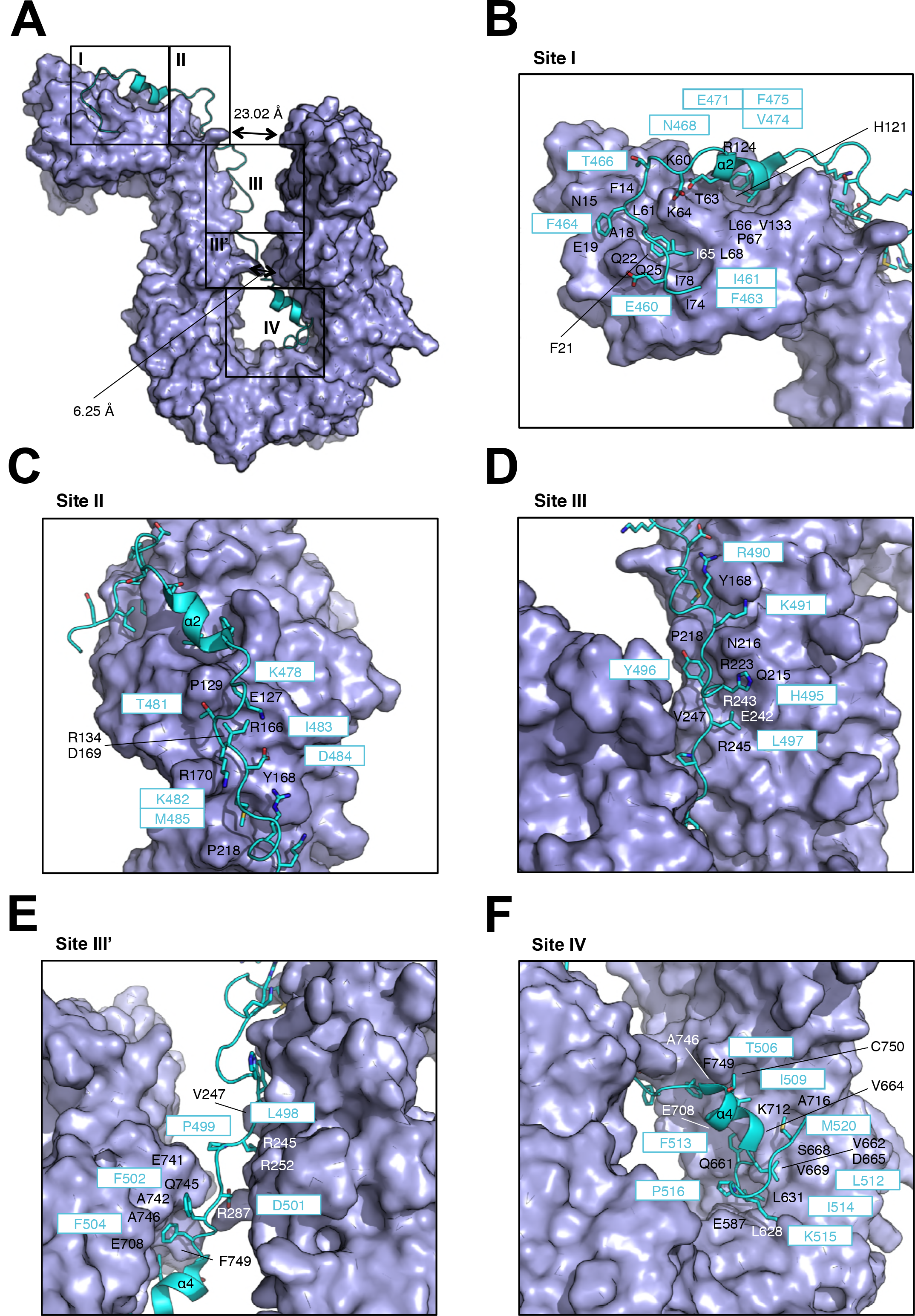
Structural details of the interaction between YCG1 and BRN1. **(A)** The molecular surface of YCG1 is shown in purple color. BRN1 is shown as light blue ribbon model. The five major contact sites (I, II, III, III’, and IV) are boxed. Representative structure was generated to use the b- and d-molecules from the reported structure of the YCG1-BRN1 subcomplex (PDB ID: 5OQQ). **(B-F)** Zoomed-in views of site I-IV. Residues of YCG1 and BRN1 are labeled in white or black and light blue, respectively.

## REFERENCES

Adams, P. D., Afonine, P. V., Bunkóczi, G., Chen, V. B., Davis, I. W., Echols, N., … Zwart, P. H. (2010). PHENIX: A comprehensive Python-based system for macromolecular structure solution. Acta Crystallographica Section D: Biological Crystallography, 66(2), 213–221. https://doi.org/10.1107/S0907444909052925

Cuylen, S., Metz, J., & Haering, C. H. (2011). Condensin structures chromosomal DNA through topological links. Nature Structural and Molecular Biology, 18(8), 894–901. https://doi.org/10.1038/nsmb.2087

Emsley, P., & Cowtan, K. (2004). Coot: Model-building tools for molecular graphics. Acta Crystallographica Section D: Biological Crystallography, 60(12 I), 2126–2132. https://doi.org/10.1107/S0907444904019158

Evans, P. (2006). Scaling and assessment of data quality. Acta Crystallographica Section D: Biological Crystallography, 62(1), 72–82. https://doi.org/10.1107/S0907444905036693

Ganji, M., Shaltiel, I. A., Bisht, S., Kim, E., Kalichava, A., Haering C. H., Dekker, C. (2018). Real-time imaging of DNA loop extrusion by condensin. Science, 360(6384), 102–105.

Griese, J. J., Witte, G., & Hopfner, K. P. (2010). Structure and DNA binding activity of the mouse condensin hinge domain highlight common and diverse features of SMC proteins. Nucleic Acids Research, 38(10), 3454–3465. https://doi.org/10.1093/nar/gkq038

Haering, C.H., Schoffnegger, D., Nishino, T., Helmhart, W., Nasmyth, K., & Lowe, J. (2004). Structure and stability of cohesin’s Smc1-kleisin interaction. Mol. Cell, 15(6), 951–964.

Hagstrom, K. a, Hagstrom, K. a, Holmes, V. F., Holmes, V. F., Cozzarelli, N. R., Cozzarelli, N. R., … Meyer, B. J. (2002). C. elegans. Genes & Development, 729–742. https://doi.org/10.1101/gad.968302.and

Hara, K., Zheng, G., Qu, Q., Liu, H., Ouyang, Z., Chen, Z., … Yu, H. (2014). Structure of cohesin subcomplex pinpoints direct shugoshin-Wapl antagonism in centromeric cohesion. Nature Structural and Molecular Biology, 21(10), 864–870. https://doi.org/10.1038/nsmb.2880

Hirano, T. (2016). Condensin-Based Chromosome Organization from Bacteria to Vertebrates. Cell, 164(5), 847–857. https://doi.org/10.1016/j.cell.2016.01.033

Ishida, T., & Kinoshita, K. (2007). PrDOS: Prediction of disordered protein regions from amino acid sequence. Nucleic Acids Research, 35(SUPPL.2), 460–464. https://doi.org/10.1093/nar/gkm363

Ivanov, D., & Nasmyth, K. (2005). A topological interaction between cohesin rings and a circular minichromosome. Cell, 122(6), 849–860. https://doi.org/10.1016/j.cell.2005.07.018

Ivanovic, T., Choi, J. L., Whelan, S. P., van Oijen, A. M., & Harrison, S. C. (2013). Influenza-virus membrane fusion by cooperative fold-back of stochastically induced hemagglutinin intermediates. eLife, 2013(2), 1–20. https://doi.org/10.7554/eLife.00333

Kabsch, W. (2010). Xds. Acta Crystallographica Section D: Biological Crystallography, 66(2), 125–132. https://doi.org/10.1107/S0907444909047337

Kikuchi, S., Borek, D. M., Otwinowski, Z., Tomchick, D. R., & Yu, H. (2016). Crystal structure of the cohesin loader Scc2 and insight into cohesinopathy. Proceedings of the National Academy of Sciences, 113(44), 12444–12449. https://doi.org/10.1073/pnas.1611333113

Kimura, K., & Hirano, T. (1997). ATP-dependent positive supercoiling of DNA by 13S condensin: A biochemical implication for chromosome condensation. Cell, 90(4), 625–634. https://doi.org/10.1016/S0092-8674(00)80524-3

Kinoshita, K., Kobayashi, T. J., & Hirano, T. (2015). Balancing acts of two HEAT subunits of condensin I support dynamic assembly of chromosome axes. Developmental Cell, 33(1), 94–107. https://doi.org/10.1016/j.devcel.2015.01.034

Kschonsak, M., Merkel, F., Bisht, S., Metz, J., Rybin, V., Hassler, M., & Haering, C. H. (2017). Structural Basis for a Safety-Belt Mechanism That Anchors Condensin to Chromosomes. Cell, 171(3), 588–600.e24. https://doi.org/10.1016/j.cell.2017.09.008

Martin, C. A., Murray, J. E., Carroll, P., Leitch, A., Mackenzie, K. J., Halachev, M., … Jackson, A. P. (2016). Mutations in genes encoding condensin complex proteins cause microcephaly through decatenation failure at mitosis. Genes and Development, 30(19), 2158–2172. https://doi.org/10.1101/gad.286351.116

Neuwald, A. F., & Hirano, T. (2000). HEAT repeats associated with condensins, cohesins, and other complexes involved in chromosome-related functions. Genome Research, 10(10), 1445–1452. https://doi.org/10.1101/gr.147400

Onn, I., Aono, N., Hirano, M., & Hirano, T. (2007). Reconstitution and subunit geometry of human condensin complexes. EMBO Journal, 26(4), 1024–1034. https://doi.org/10.1038/sj.emboj.7601562

Ouyang, Z., Zheng, G., Tomchick, D. R., Luo, X., & Yu, H. (2016). Structural Basis and IP6 Requirement for Pds5-Dependent Cohesin Dynamics. Molecular Cell, 62(2), 248–259. https://doi.org/10.1016/j.molcel.2016.02.033

Piazza, I., Rutkowska, A., Ori, A., Walczak, M., Metz, J., Pelechano, V., … Haering, C. H. (2014). Association of condensin with chromosomes depends on DNA binding by its HEAT-repeat subunits. Nature Structural and Molecular Biology, 21(6), 560–568. https://doi.org/10.1038/nsmb.2831

Strick, T., Kawaguchi, T., & Hirano, T. (2004). Real-time detection of single-molecule DNA compaction by condensin I. Current Biology, 14, 874–880.

St-Pierre, J., Douziech, M., Bazile, F., Pascariu, M., Bonneil, É., Sauvé, V., … D’Amours, D. (2009). Polo Kinase Regulates Mitotic Chromosome Condensation by Hyperactivation of Condensin DNA Supercoiling Activity. Molecular Cell, 34(4), 416–426. https://doi.org/10.1016/j.molcel.2009.04.013

Sutani, T., Sakata, T., Nakato, R., Masuda, K., Ishibashi, M., Yamashita, D., … Shirahige, K. (2015). Condensin targets and reduces unwound DNA structures associated with transcription in mitotic chromosome condensation. Nature Communications, 6, 1–13. https://doi.org/10.1038/ncomms8815

Terakawa, T., Bisht, S., Eeftens, J. M., Dekker, C., Haering, C. H., Greene, E. C. (2017). The condensin complex is a mechanochemical motor that translocates along DNA. Science, 358(6363), 672–676.

Uhlmann, F. (2016). SMC complexes: from DNA to chromosomes. Nature Reviews Molecular Cell Biology, 17, 399–412.

Yoshimura, S. H., & Hirano, T. (2016). HEAT repeats–versatile arrays of amphiphilic helices working in crowded environments? Journal of Cell Science, 129(21), 3963–3970. http://jcs.biologists.org/content/129/21/3963.long

